# Independent Filter Analysis for Group Discrimination in fMRI

**DOI:** 10.1101/2025.11.20.689235

**Authors:** Zain Souweidane, Alberto Llera, Stephen M. Smith, Christian F. Beckmann

**Affiliations:** Donders Institute for Brain, Cognition and Behaviour, Radboud University, Nijmegen, the Netherlands; Department for Cognitive Neuroscience, Radboud University Medical Centre Nijmegen, Nijmegen, the Netherlands; R&D Group, LIS Data Solutions, Santander, Spain; Oxford Centre for Functional MRI of the Brain, University of Oxford, Oxford, United Kingdom

**Keywords:** Group-level fMRI Analysis, Supervised Dimensionality Reduction, Functional Connectivity, Spatial Filtering, Independent Component Analysis (ICA), Independent Filter Analysis (IFA)

## Abstract

Traditional group-level fMRI analysis approaches, such as Independent Component Analysis (ICA), often rely on unsupervised dimensionality reduction to map subjects into a common feature space. While effective for capturing common variance across all subjects, the preservation of discriminative features between groups of participants is not guaranteed. To address this limitation, we introduce Independent Filter Analysis (IFA), a supervised extension of group ICA that explicitly models group-discriminative information as part of the dimensionality reduction steps. Prior to unmixing, IFA constructs a subspace that simultaneously retains both shared and group-specific information, enhancing sensitivity to group effects while preserving biological interpretability. We validated IFA using simulated data and paired condition comparisons from three Human Connectome Project (HCP) tasks. In the simulation, IFA achieved 95% classification accuracy, outperforming group ICA, which failed to detect subtle group differences. On the HCP data, IFA increased network matrix classification accuracy by up to 15% and produced spatial maps that more precisely reflected task-relevant differences.

## 1 Introduction

Group-level analysis of functional MRI (fMRI) data has become an established approach for investigating functional networks across populations (Dadi et al., 2019; Finn & Rosenberg, 2021; Smith et al., 2015). Individuals’ fMRI data are mapped to a common space using a shared template (e.g., a set of canonical resting-state networks, RSNs) to establish correspondence across subjects (Beckmann et al., 2009; Beckmann et al., 2005; Calhoun et al., 2001; Smith et al., 2011; Varoquaux, Sadaghiani, et al., 2010). The resulting subject-level representations can then be compared to identify patterns of functional connectivity that are shared across individuals or—more typically—differ between subgroups such as patients vs control participants (Damoiseaux et al., 2006; Filippini et al., 2009; Greicius et al., 2004; Rabany et al., 2019; Varoquaux, Baronnet, et al., 2010; Whitfield-Gabrieli et al., 2009). One fundamental limitation of these approaches is that they constrain subject-level variability to the structure defined by the shared template, potentially overlooking clinically relevant characteristics of individuals (Bijsterbosch et al., 2018; Smith, 2012). This limitation arises from the inherent tension in creating the initial shared subspace, which must balance consistency across subjects while preserving relevant inter-individual differences.

Overcoming this limitation requires establishing a group-level template that explicitly models components that support the downstream task, such as classification, regression, or subgroup discovery. To understand the relationship between two subgroups, the relevant components include the networks shared by all subjects, the within-group similarities that reflect subgroup homogeneity, and the between-group differences that distinguish the subgroups. Traditional group-level methods, such as Independent Component Analysis (ICA), model the first aspect while largely ignoring the latter two. To address this gap, we propose Independent Filter Analysis (IFA), a supervised extension of ICA that captures both shared and group-discriminative features.

Group-level ICA is biased toward capturing shared signals due to the use of Principal Component Analysis (PCA) to reduce the dimensionality of the data (Beckmann & Smith, 2004; Sui et al., 2010). PCA is performed on a temporally concatenated dataset to produce a lower-dimensional representation that emphasizes consistency across subjects rather than variability between them (Calhoun et al., 2001; Smith et al., 2014; Varoquaux, Sadaghiani, et al., 2010). As a result, group differences are only preserved if they align with the group-average-based components (Chang, 1983). In practice, this alignment is not guaranteed. Clinically meaningful group effects can occur in directions of lower variance (Zhao et al., 2022). Additionally, the subspace can be biased toward capturing variance driven predominantly by the group with more subjects or a cleaner signal.

To address this limitation, supervised dimensionality reduction has been widely adopted (Foley & Sammon, 1975; Fukunaga & Koontz, 1970; Ghojogh et al., 2023; Sheng Zhang & Sim, 2006). These approaches prioritize group-separating directions over directions of maximal variance. In fMRI analysis, related techniques have been applied to increase downstream classification accuracy (Andersen et al., 2012; Eavani et al., 2014; Zhang et al., 2020). More recent work aimed at modeling inter-group relationships has further focused on decompositions that preserve both shared and group-specific features (Ghayem et al., 2023; Jin et al., 2023). However, most supervised ICA pipelines incorporate class labels only during the unmixing stage (Du et al., 2023; Maneshi et al., 2016; Tabas et al., 2014; Vahdat et al., 2012) or rely on PCA-based dimensionality reduction limited to task-derived features (Sui et al., 2009). We developed IFA to overcome these methodological limitations by incorporating group-level discrimination into the dimensionality reduction process.

IFA introduces a novel framework for group-level fMRI analysis that jointly preserves shared and group-discriminative information. The shared subspace captures canonical functional networks and structured noise. This provides context for interpreting group differences and ensures discriminative signals remain distinct from nuisance variance. Directly modeling discriminative information prevents subtle but meaningful group-specific effects from being discarded during dimensionality reduction.

We validated IFA on both a controlled simulation and paired comparisons within three tasks from the Human Connectome Project (HCP). These experiments demonstrate that IFA improves the retention of group-specific information while preserving biological interpretability. In the simulated benchmark where group-specific effects were expressed in low-variance directions, IFA achieved 95% classification accuracy of the resulting network matrices, whereas group ICA (GICA) (Nickerson et al., 2017) failed to retain sufficient discriminant information. On the HCP data, IFA improved network matrix classification accuracy by 10–15%. Additionally, the IFA derived spatial maps contained more discriminative power on average and captured focal task-relevant differences, such as the fusiform face area, that GICA largely missed.

## 2 Methods

IFA is a supervised extension of group ICA (GICA) designed to extract spatial components that capture common patterns across subjects while highlighting differences between groups. Like traditional GICA, IFA performs dimensionality reduction prior to ICA unmixing. It differs by modifying the reduced component space to maximize group differences.

For each subject *j* = 1, …, *n*, let **X**^(*j*)^ ∈ ℝ^*t*×*s*^ represent the data matrix, where *n* is the number of subjects, *t* the number of time-points, and *s* the number of grayordinates (voxels and/or vertices). GICA begins by temporally concatenating subjects to form the full data matrix **X** ∈ ℝ^(*n*·*t*)×*s*^. This matrix is then factorized using singular value decomposition (SVD):

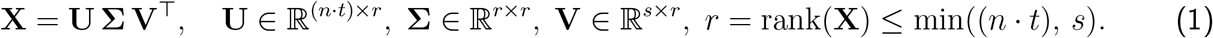

Here, **U** contains the left singular vectors, **Σ** the singular values, and **V** the spatial eigenvectors. The first *q* ≤ *r* columns of **V, V**_*q*_ = **V**_[:,1:*q*]_ ∈ ℝ^*s*×*q*^, are retained to form the component space that maximizes explained variance (i.e., PCA). These components are rotated by the unmixing matrix **Q** ∈ St(*q, q*), estimated using FastICA (Hyvärinen & Oja, 2000):

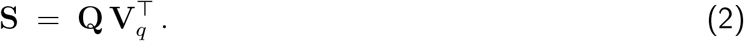

This results in statistically independent spatial maps, **S** ∈ ℝ^*q*×*s*^, that serve as a group template.

IFA extends this framework by combining the variance-explaining subspace **V**_*q*_ with a discriminant subspace **D** ∈ ℝ^*s*×*l*^, derived via supervised dimensionality reduction. As discussed in Section 2.1, **V**_*q*_ is orthogonal to **D**, allowing the full signal space to be obtained by concatenating the two subspaces. The augmented component space is unmixed by **Q** ∈ St(*q* + *l, q* + *l*):

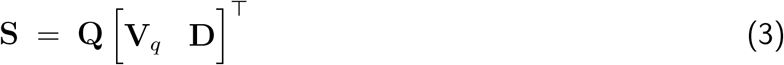

yielding the final spatial maps **S** ∈ ℝ^(*q*+*l*)×*s*^.

IFA ensures that the group-level template, **S**, captures both shared variance and group-level differences. **S** is then used in dual regression (Nickerson et al., 2017) to derive the subject-level representations. The full IFA pipeline is described in Section 2.1, followed by Section 2.2, which details the construction of the discriminant subspace and compares two implementations. One models subject-level variability in parcellated space, while the other operates directly in grayordinate space using group-averaged representations. Figure 1 provides a high-level overview of the pipeline.

**Figure 1:**
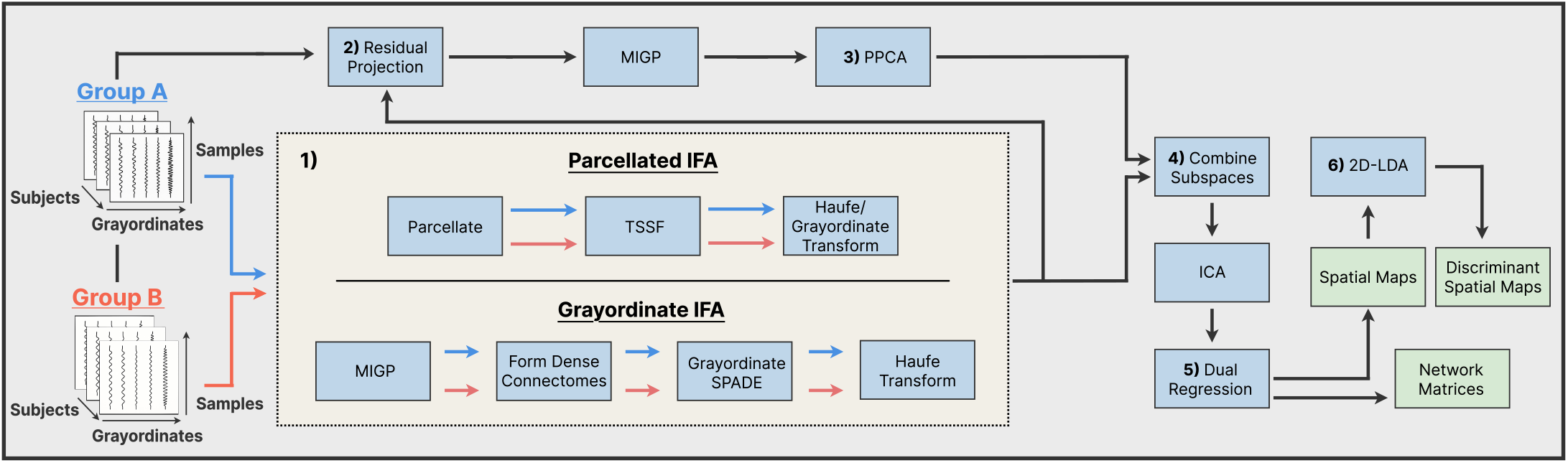
Independent Filter Analysis (IFA) pipeline. **1)** Discriminant filters are learned through one of the two complementary branches, Parcellated IFA (top) or the Grayordinate IFA (bottom). **2)** Each subject is projected onto the residual space orthogonal to these discriminant filters. **3)** MIGP + PPCA extract components that capture variance shared by both groups. **4)** The shared PCA basis and the discriminant bases are concatenated and unmixed with FastICA. **5)** Dual regression produces subject-level spatial maps and time-courses, from which network matrices are formed. **6)** 2D-LDA is applied to the subject-level spatial maps to extract spatial directions that maximize between-group separation, consolidating discriminant information.

### 2.1 Pipeline Overview

IFA begins by constructing a discriminative subspace, **D**, through a supervised dimensionality reduction step based on the Fukunaga–Koontz Transform (Fukunaga & Koontz, 1970). This step can be implemented in either parcellated or grayordinate space, referred to as Parcellated IFA and Grayordinate IFA, respectively. In Parcellated IFA, filters are derived from subject-level parcellated covariance matrices and then projected back to grayordinate space. In Grayordinate IFA, filters are estimated directly in grayordinate space from group-average covariance matrices. Starting at Step 2 (Figure 1), both pipelines are identical and operate entirely in grayordinate space. Details of the two variants are provided in Section 2.2, while the remainder of this section outlines the subsequent, shared steps.

To isolate variance not captured by the discriminative subspace, each subject’s data **X**^(*j*)^ is projected onto the residual space orthogonal to **D**:

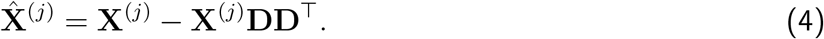

Residual subject matrices are temporally concatenated to form the full residual data matrix 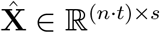.

Unsupervised dimensionality reduction is applied to the residualized data to estimate the variance-explaining subspace^1^. Since 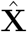 is often too large to decompose directly, MELODIC’s Incremental Group-PCA (MIGP) is first applied to reduce the number of samples (Smith et al., 2014). This is followed by Probabilistic Principal Component Analysis (PPCA) (Beckmann & Smith, 2004) to estimate **V**_*q*_ ∈ ℝ^*s*×*q*^. By construction, 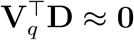. The two subspaces are concatenated and unmixed using FastICA (Eq. 3), yielding group-level spatial maps **S** ∈ ℝ^(*q*+*l*)×*s*^.

Final subject-specific representations are derived from the group spatial maps using dual regression (Nickerson et al., 2017). Subject-level network matrices are formed from the covariance of component time courses estimated in the first stage, with regularization applied using Oracle Approximating Shrinkage (OAS) (Chen et al., 2010). The second stage typically involves regressing the subject’s data onto the normalized time courses, which are often highly correlated. To improve the reliability of the reconstructed maps, we replaced the standard second-stage regression with ElasticNet (Zou & Hastie, 2005). There, the *ℓ*_2,2_ penalty mitigates sensitivity to correlated predictors and noise, while the *ℓ*_1,1_ penalty favors sparseness. This regularization reflects the assumption that only a subset of components contributes to each grayordinate. Details of hyperparameter selection are provided in Appendix A.

In addition to subject-level spatial maps and network matrices, we derive a complementary set of discriminant spatial maps for each subject to better isolate group differences. Although the discriminant components are orthogonal to the shared variance components, they may still share higher-order statistical dependencies. As a result, ICA unmixing distributes discriminative information across spatial maps in a way that embeds it within functional networks and maximizes spatial independence. Consequently, no single spatial map is guaranteed to align with the spatial direction that maximizes the separation between groups.

To identify linear combinations of spatial maps that maximize between-group separation, we apply two-dimensional Linear Discriminant Analysis (2D-LDA) (Ye et al., 2004) to the subject-level maps. 2D-LDA constructs a transformation matrix, **H**, which defines the linear combination of spatial maps that maximizes group separation. Each subject’s spatial representation, **S**^(*j*)^, is projected onto the discriminant directions defined by **H**, resulting in a transformed representation 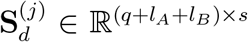. Since the total discriminant information within the subspace is preserved, the transformation redistributes discriminant information across the maps, concentrating it within the top components. The rows of 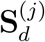 are ordered by decreasing discriminative power, with the top capturing directions of greatest group separation and the bottom reflecting the least. This representation highlights regions of group difference independent of their functional roles. A complete derivation of this discriminant transformation is provided in Appendix B.

### 2.2 Discriminant Subspace

The discriminant subspace should reduce the dimensionality of the data while preserving meaningful group differences. Spatial Patterns for Discriminant Estimation (SPADE), an application of FKT in fMRI data, has proven to be effective at isolating group differences using a small set of spatial filters (Llera et al., 2020). These filters are derived to maximize variance in the group-mean covariance of one class, 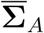, while minimizing it in the other, 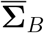. This is achieved by solving a generalized eigenvalue decomposition (GEVD) which can equivalently be reformulated as a generalized Rayleigh quotient:

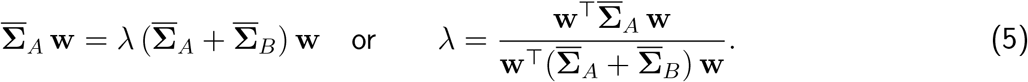

Eigenvectors, **w**, with corresponding eigenvalues, *λ* ∈ [0, 1], near 1 or 0 indicate directions of dominant variance in Group A or Group B, respectively. Filters with eigenvalues near 0.5 are non-discriminative and typically discarded. The eigenvectors optimally separate samples drawn from the class-average distributions (Huo et al., 2003), which maximizes the separation between the group-average covariances (Barachant et al., 2010). Ghojogh et al. (2019) and Cohen (2022) provide detailed discussions on the GEVD and its application in neuroimaging.

Both IFA variants use different extensions of this FKT framework. In both cases, for each group *g* ∈ {A, B}, this yields a set of spatial filters 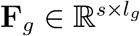, whose *l*_*g*_ columns define directions of group-specific variance. Ideally, **F**_*g*_ would capture patterns reflecting meaningful signal unique to Group *g*. However, since Eq. 5 defines a discriminative model, the resulting patterns may reflect model effects such as overfitting to correlated noise, scalings due to regularization parameters, or suppression of variance associated with the opposite group. To remove these model dependencies and map filter weights to brain function, **F**_*g*_ is Haufe-transformed (Haufe et al., 2014) using the group-specific data, yielding 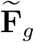. These filters are concatenated to form the full discriminative subspace 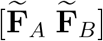. The resulting filters are orthonormal only with respect to the joint covariance (Cohen, 2022). A QR decomposition is applied to obtain a basis that is orthonormal in the data space while spanning the same discriminative subspace, yielding 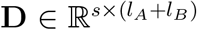. These steps define the general formulation of the discriminant subspace used in IFA. The exact details of the subspace construction for Parcellated IFA and Grayordinate IFA are described below.

*Parcellated IFA* — Because SPADE bases group discrimination entirely on class-mean covariances, it is less effective when within-class heterogeneity is high. To address this limitation, we apply a variant of Tangent Space Spatial Filtering (TSSF) (Xu et al., 2019), originally developed for EEG analysis, to parcellated fMRI data. The spatial filters estimated by TSSF reconstruct the direction that maximizes between-group separation on the Riemannian manifold of covariance matrices. Subject-level covariance matrices are first mapped to a vector space by projecting them onto the tangent plane centered at the Fréchet mean across both groups. A linear classifier is then trained on these vectorized covariances, and its weight vector is reshaped into a symmetric matrix used to extract spatial filters via an EVD. Further details on TSSF, its connection to SPADE, and adaptations under the Log-Euclidean metric are provided in Appendix C.

Since training a linear classifier on dense subject-level connectomes is computationally intractable, covariance matrices are instead computed from parcellated data. The parcellated covariances are regularized using OAS, which shrinks the empirical covariance toward a scaled identity matrix to improve conditioning and ensure positive definiteness. TSSF is applied to these matrices, yielding discriminant filters 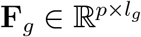, where *p* is the number of parcels. Dual regression is used to project each filter from parcel to grayordinate space, which also functions as the Haufe transform, producing 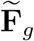 (implementation details in Appendix D).

*Grayordinate IFA* — Although the application of TSSF effectively identifies filters that capture subject-level variability, its reliance on parcellated representations may obscure finer group differences present in the full grayordinate space. However, extending TSSF directly to subject-level grayordinate connectomes is infeasible, since a single dense connectome can contain billions of elements. Grayordinate IFA therefore trades within-class subject-level heterogeneity for increased spatial resolution by using the group-average dense connectome as a summarized representation of each group. To accomplish this, we extend SPADE to grayordinate-level data by applying it to group-average dense connectomes, **C**_*A*_ and **C**_*B*_ ∈ ℝ^*s*×*s*^. The connectomes are formed from the output of MIGP applied to each group separately and regularized with OAS. Discriminant filters are then derived by solving Equation 5 with **C**_*A*_ and **C**_*B*_ in place of 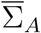 and 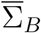, respectively.

To perform this computation efficiently in the high-dimensional setting, we use Locally Optimal Block Preconditioned Conjugate Gradient Descent (LOBPCG) (Knyazev, 2001), an iterative solver that avoids matrix inversion and supports preconditioning to accelerate convergence. The resulting eigenvectors define spatial filters 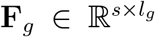 that emphasize group-specific variance while preserving full spatial resolution. Since these filters already lie in grayordinate space, dual regression is not required. Instead, we apply the Haufe-transform directly to each group’s filters using their corresponding dense connectome:

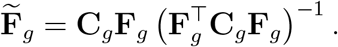

This yields interpretable group-discriminative patterns without the spatial constraints imposed by parcellation. While the original SPADE formulation implicitly assumes a Euclidean metric for group-average formation, its discriminant distance estimation naturally reflects the geometry induced by the Riemannian metric (Barachant et al., 2010). As an alternative formulation, we derive a Log-Euclidean variant (Arsigny et al., 2007) of Grayordinate SPADE in Appendix E, where the Log-Euclidean metric is applied to both mean estimation and discriminant distance computation. This formulation preserves key matrix properties while offering numerical stability and computational efficiency (Chevallier et al., 2021).

In practice, for both SPADE and TSSF, only a few of the top and bottom filters are retained, since these capture the majority of meaningful group differences (Xu et al., 2019; Zhang et al., 2020). The eigenvalues can be transformed, squared in the case of TSSF or 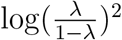 for SPADE (Barachant et al., 2010), and visualized in a scree plot to aid in filter selection. For automatic model order selection, the transformed eigenvalues can be compared against a null distribution generated via permutation testing. When using TSSF, the Haufe transform should be applied to the classifier weight vector in the tangent space before the eigenvalue decomposition (EVD) to ensure that the resulting eigenvalues reflect genuine group-level effects rather than artifacts of the classifier, enabling valid permutation testing.

### 2.3 Evaluation

We evaluated the performance of IFA, both Parcellated IFA and Grayordinate IFA, against standard group ICA (GICA) on simulated data, reported in 3.4, and task-based fMRI data from the Human Connectome Project (HCP). For the HCP analyses, each pipeline was applied to preprocessed CIFTI data (Glasser et al., 2013) from three paired-condition tasks: Face vs. Tools (Working Memory), Relational vs. Match (Relational), and Reward vs. Punishment (Incentive Processing). Classes were defined by contrasting conditions completed by the same subjects, creating a within-subject (paired) comparison. These conditions were chosen for their low effect sizes (Barch et al., 2013), which present a realistic and challenging benchmark for group-level discriminative analysis.

In Parcellated IFA, supervised dimensionality reduction was performed in the parcellated space defined by the Multi-Modal Parcellation (MMP) atlas (Glasser et al., 2016). For Riemannian-based analyses, including TSSF and network matrix analysis, the Log-Euclidean metric was used based on prior evidence of computational efficiency and robust performance (Chevallier et al., 2021; Pervaiz et al., 2020). To evaluate generalization, all pipelines were assessed using 5-fold cross-validation stratified by class label and family ID, ensuring that members of the same family were assigned entirely to either the training or test set. Test folds were excluded from all stages of network matrix and spatial map generation, as well as from hyperparameter tuning. The simulated dataset included 100 subjects per class, while each HCP task involved approximately 1,000 paired samples, resulting in 1,000 samples per class.

Discriminative performance was compared across network matrices (Section 3.1), spatial maps, and discriminant spatial maps (Section 3.2) generated by each pipeline. A fixed model order of 10 components was used for all pipelines, selected to balance interpretability with the ability to capture task-relevant networks. Both IFA variants included six variance-explaining components and four discriminant components. For traditional group ICA, we matched this dimensionality using 10 variance-explaining components. However, because performance is sensitive to model order, we provide a detailed comparison across multiple orders in Section 3.3. We reran the network matrix classification analysis using model orders of 5, 10, 15, 25, 35, and 50. For each model order, IFA matched the total dimensionality of GICA by replacing four variance-explaining components with four discriminant components.

To evaluate the extent to which the resulting network matrices retained group discriminant information, each network matrix was first projected to the tangent space centered at the Riemannian mean of the training matrices. Group differences were then evaluated using SVM classification (C = 0.1) (Pervaiz et al., 2020) and permutation-based t-tests with maximum statistic correction (*α* = 0.05 and 10,000 permutations) (Varoquaux, Baronnet, et al., 2010). To visualize group separation, we applied TSSF to the network matrices, extracting two discriminative filters. Each subject’s demeaned first-stage time series was projected onto these filters, and the log-variance of the projected signals was computed. We trained a linear classifier on the log-variance features and measured the Euclidean distance between group means as an approximation for their Riemannian separation (Barachant et al., 2010; Xu et al., 2019).

Similar to the network matrix analysis, we evaluated the spatial maps from all pipelines using per-map linear SVM classification (*C* = 0.1) and vertex-wise paired *t*-tests of activation differences across conditions. For classification, individual subject-level spatial maps were used as features. Vertex-wise permutation-based *t*-tests were conducted with maximum-statistic correction (*α* = 0.05 and 10,000 permutations). A summary map was generated by taking the minimum *p*-value at each vertex across components to summarize results. These analyses were repeated on the 2D-LDA transformed spatial maps.

In addition to testing performance on HCP data, we generated two simulated datasets to provide a controlled benchmark. All pipelines were tested on the simulated datasets using a model order of 10, with two discriminant components specified for IFA. For both datasets, we simulated two classes each containing 200 subjects, where subjects’ data matrices were 200 time points by 91, 282 vertices. Each subject was formed from the combination of a fixed rank 30 matrix representing shared components, a rank two matrix defining the discriminant subspace, and a rank 168 subject-specific isotropic noise matrix.

For both simulated datasets, we embedded the two–dimensional discriminant subspace after the 10th eigenvalue to model a scenario where discriminant information is not represented in the leading eigenspace. Within this subspace, the proportion of variance each eigenvector explained for a given subject was drawn from a class-specific Beta distribution. In Dataset 1, each eigenvector predominantly explained variance in one class but not the other, with low within-class spread per filter. Dataset 2 preserved the opposing discriminant eigenvectors structure, but changed the within-class spread such that one eigenvector had low spread while the other had high spread. Figure 8 visualizes the placement of the discriminant subspace in the group-level eigenspectrum as well as the two-dimensional discriminant structures from both datasets.

## 3 Results

### 3.1 Network Matrix Analysis

Across all tasks, both IFA variants outperformed GICA in tangent-space classification accuracy (Table 1). Relative to ICA, IFA improved accuracy by approximately 10–15% for Face vs. Tools and Relational vs. Match, and by 5–10% for Reward vs. Punishment. For Face vs. Tools and Relational vs. Match, the two IFA variants performed comparably. Grayordinate IFA achieved slightly higher accuracies, potentially reflecting its ability to capture subtle spatial differences at the grayordinate level. However, for Reward vs. Punishment, which includes high within-group variability and weaker task contrasts (Barch et al., 2013), Parcellated IFA outperformed Grayordinate IFA. Grayordinate IFA’s reduced performance may reflect its susceptibility to overfitting in noisy or ill-conditioned settings, caused by its reliance on high-dimensional features and group-averaged representations. This is further conveyed by its higher standard deviation across folds for the Relational vs. Match and Reward vs. Punishment tasks.

**Table 1:**
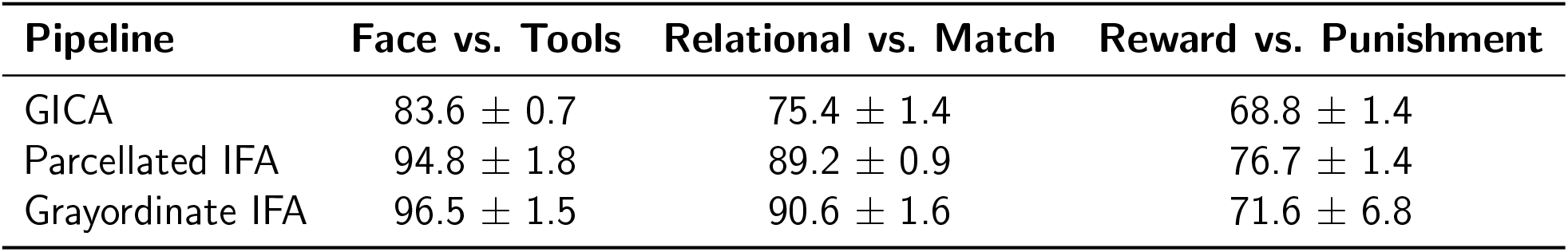
Tangent Network Matrix Classification Accuracy (%, ± SD) Across Folds.

| Pipeline | Face vs. Tools | Relational vs. Match | Reward vs. Punishment |
| --- | --- | --- | --- |
| GICA | 83.6 ± 0.7 | 75.4 ± 1.4 | 68.8 ± 1.4 |
| Parcellated IFA | 94.8 ± 1.8 | 89.2 ± 0.9 | 76.7 ± 1.4 |
| Grayordinate IFA | 96.5 ± 1.5 | 90.6 ± 1.6 | 71.6 ± 6.8 |

2D log-variance projections derived from TSSF filters yielded classification results consistent with tangent-space analysis. As shown in Figure 2, both IFA variants increased classification accuracy and between-group separation across tasks as compared with GICA. On the Working Memory task, Grayordinate IFA resulted in higher classification accuracy than Parcellated IFA (93% vs. 90%) and a larger distance between group means (0.72 vs. 0.66). A similar result was observed for the Relational task, with Grayordinate IFA reaching 92% accuracy and a distance of 0.62, compared to 87% accuracy and 0.52 for Parcellated IFA. These results suggest that Grayordinate IFA better preserves subtle spatial differences after dimensionality reduction when group effects are reliable but modest. However, in the Reward vs. Punishment comparison Grayordinate IFA achieved lower classification accuracy than Parcellated IFA (71% vs. 76%) but a comparable or slightly greater mean distance between groups (0.45 vs. 0.42). This pattern may reflect Grayordinate IFA’s stronger emphasis on separating class means, even when overall classification accuracy is reduced under noisy conditions.

**Figure 2:**
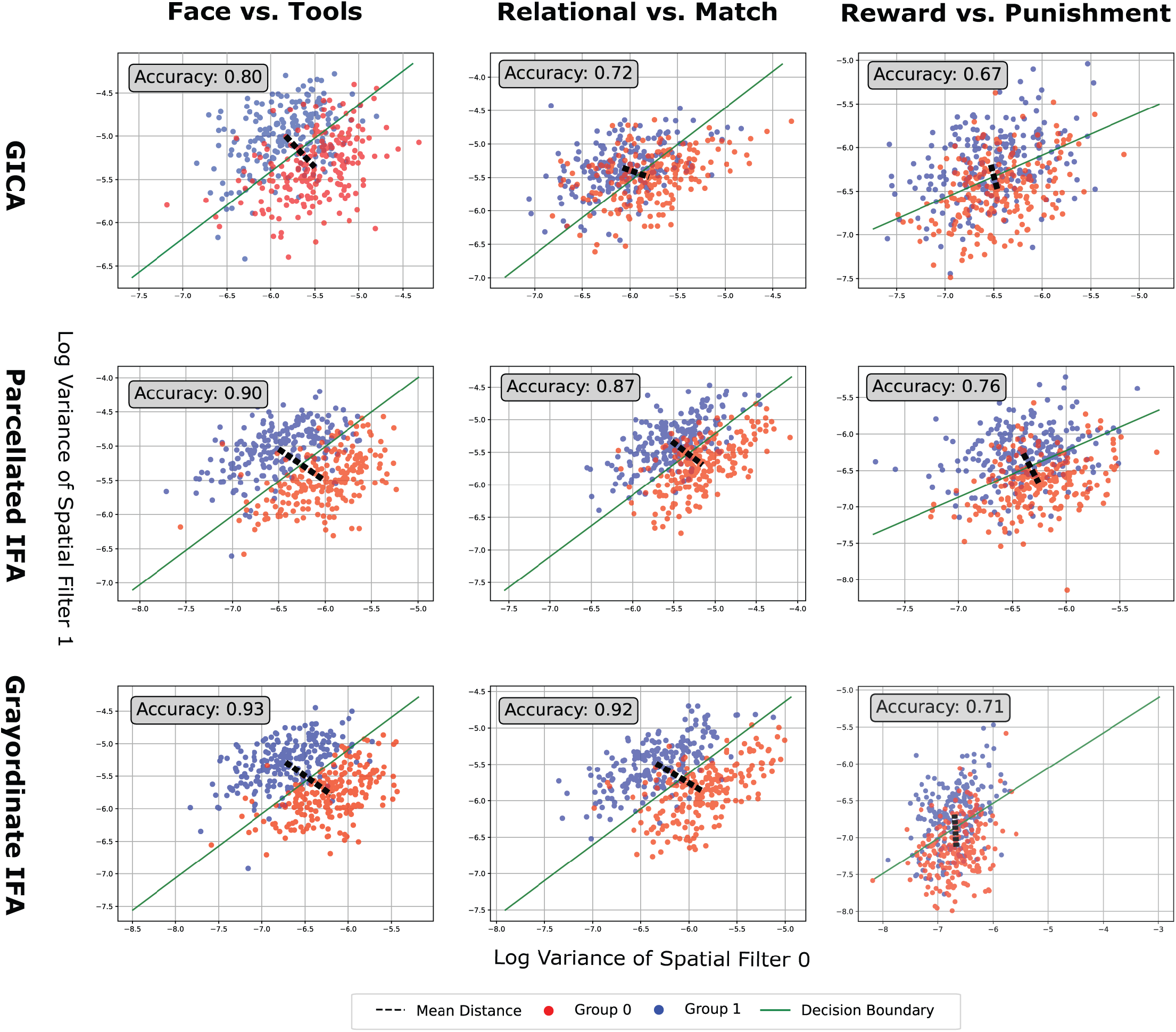
2D Log-Variance Projections Using TSSF-Trained Filters. Each plot shows the log-variance of subjects’ component time series projected into a 2D space defined by discriminant filters learned via TSSF on network matrices from each pipeline (rows) and task (columns). A linear SVM (*C* = 0.1) was trained on this 2D feature space. Points are colored by group, with a decision boundary (green) and group mean distance (dashed black line) shown. Classification accuracy is reported in each plot. These results are from a single fold, representative of the patterns observed across all folds.

The same group difference trends observed in classification accuracy are also observed in the tangent *t*-test analysis. As shown in Table 2 and Figure F, both IFA pipelines identified more significant network connections than ICA across all tasks. This emphasizes that the discriminant subspaces recovered by IFA improve both classification performance and the statistical interpretability of group-level differences.

**Table 2:** Mean Number of Significant Tangent T-Test Network Matrix Connections (out of 55 Possible) Across Folds (± SD)

| Pipeline | Face vs. Tools | Relational vs. Match | Reward vs. Punishment |
| --- | --- | --- | --- |
| GICA | 14.6 ± 2.1 | 18.6 ± 3.0 | 8.0 ± 1.6 |
| Parcellated IFA | 23.2 ± 1.1 | 23.0 ± 2.0 | 13.5 ± 1.3 |
| Grayordinate IFA | 24.4 ± 2.6 | 23.8 ± 3.9 | 12.25 ± 2.1 |

### 3.2 Spatial-Map Analysis

Results from the spatial map classification were aligned with the network matrix findings. Across all HCP task conditions, both IFA variants generally outperformed GICA in classification accuracy. The largest average gain appeared in Face vs. Tools (4.07–4.32%), followed by Relational vs. Match (2.46–2.71%), with Reward vs. Punishment showing only modest improvements. The biggest improvements occurred in maps with mid-level performance under GICA. High-performing maps changed little, as they already captured strong task-related variance, while low-performing maps showed no improvement due to lack of discriminative signal. This trend is illustrated by the inverted-U shape seen across the subplots of the left panel of Fig. 3.

**Figure 3:**
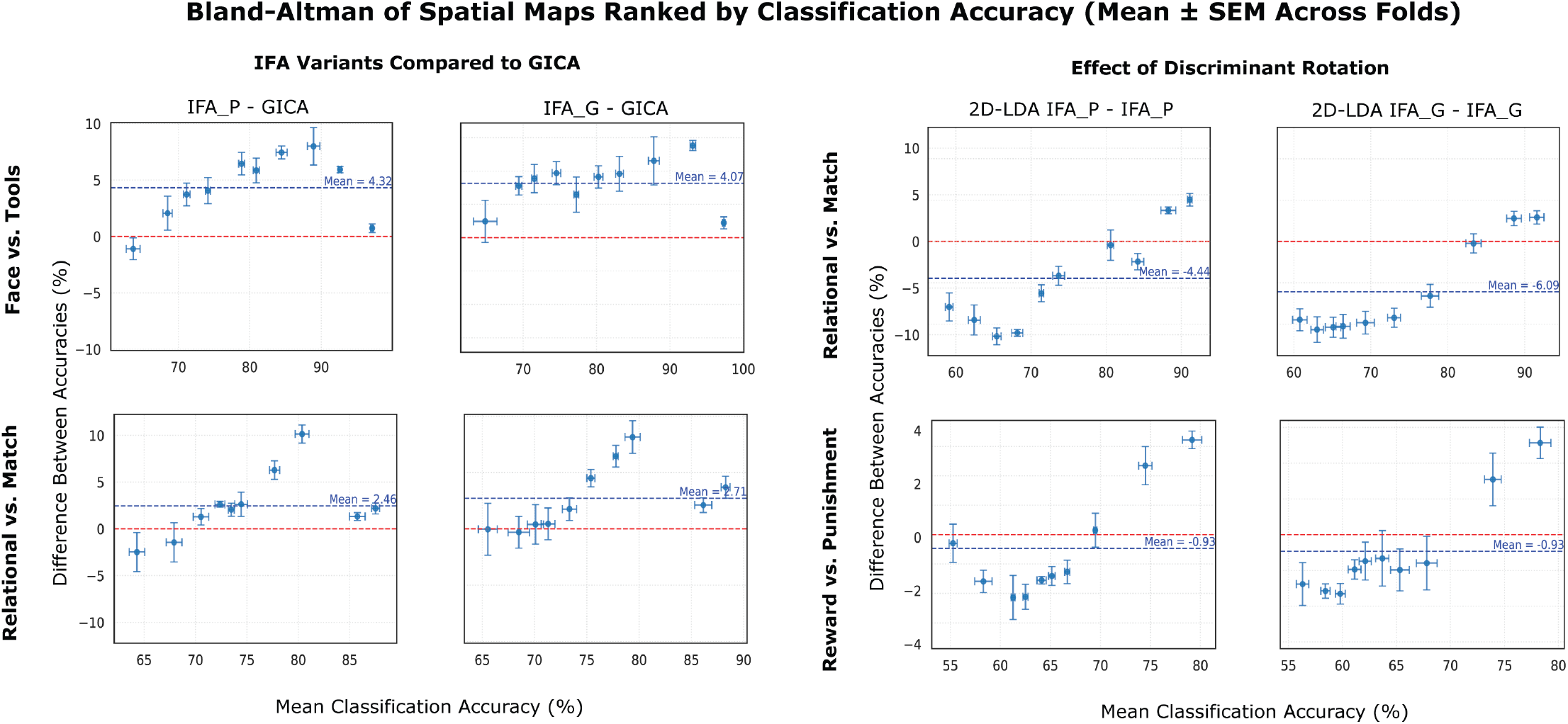
Bland-Altman analysis of ranked spatial map classification performance across pipelines. Each subplot compares two analysis pipelines based on the classification accuracy of their ranked spatial maps. For each fold, spatial maps from both pipelines were ranked by classification accuracy, and maps of the same rank were paired for comparison. Each point (ten per pipeline, matching the model order) represents the difference in accuracies between a pair (*y*-axis) plotted against their mean accuracy (*x*-axis). Values are averaged across folds, with error bars indicating the SEM. Rows represent experimental conditions and columns show different pipeline comparisons. IFA P denotes Parcellated IFA and IFA G denotes Grayordinate IFA. The left panel compares the IFA variants to GICA, and the right panel compares the 2D-LDA–rotated maps to their corresponding unrotated maps. The dashed red line at *y* = 0 denotes equal performance; positive values indicate superior performance by the first-listed method in the comparison title. This rank-wise Bland-Altman approach reveals how discriminative power is distributed across spatial map ranks and how consistently one method outperforms another across the maps.

The raw spatial map accuracies reflect how task discrimination is embedded within functional networks, whereas results from the discriminant spatial maps convey how that discriminative signal is distributed across the subspace spanned by the original components. The right panel of Fig. 3 shows how the discriminant rotation reshaped the distribution of discriminant information across maps. The 2D-LDA projection increased classification accuracy for the top-ranked maps but decreased it for lower-ranked ones. The average classification accuracy across all maps decreases, but the top two maps’ accuracies increase. This aligns with the interpretation that the projection concentrates task-related variance into a reduced set of components, while preserving the total amount of discrimination in the subspace.

The spatial *t*-tests produced findings similar to the spatial classification. In line with prior findings of Barch et al., 2013, none of the pipelines produced reliable *t*-test differences for Reward vs. Punishment. For this reason, the presented results focus on Face vs. Tools and Relational vs. Match. To assess the spatial distribution of *t*-test results, we calculated the proportion of vertices within each MMP parcel that were deemed significant based on the thresholded minimum-*p* maps from each pipeline. This allowed us to compare how each pipeline captured significant differences within a parcel, with results averaged across folds. Figure G summarizes these results for both the raw and discriminant spatial maps across the two contrasts of interest. Figure 4 shows the cortical surface visualizations from a representative fold of discriminant map comparison for Face vs. Tools.

**Figure 4:**
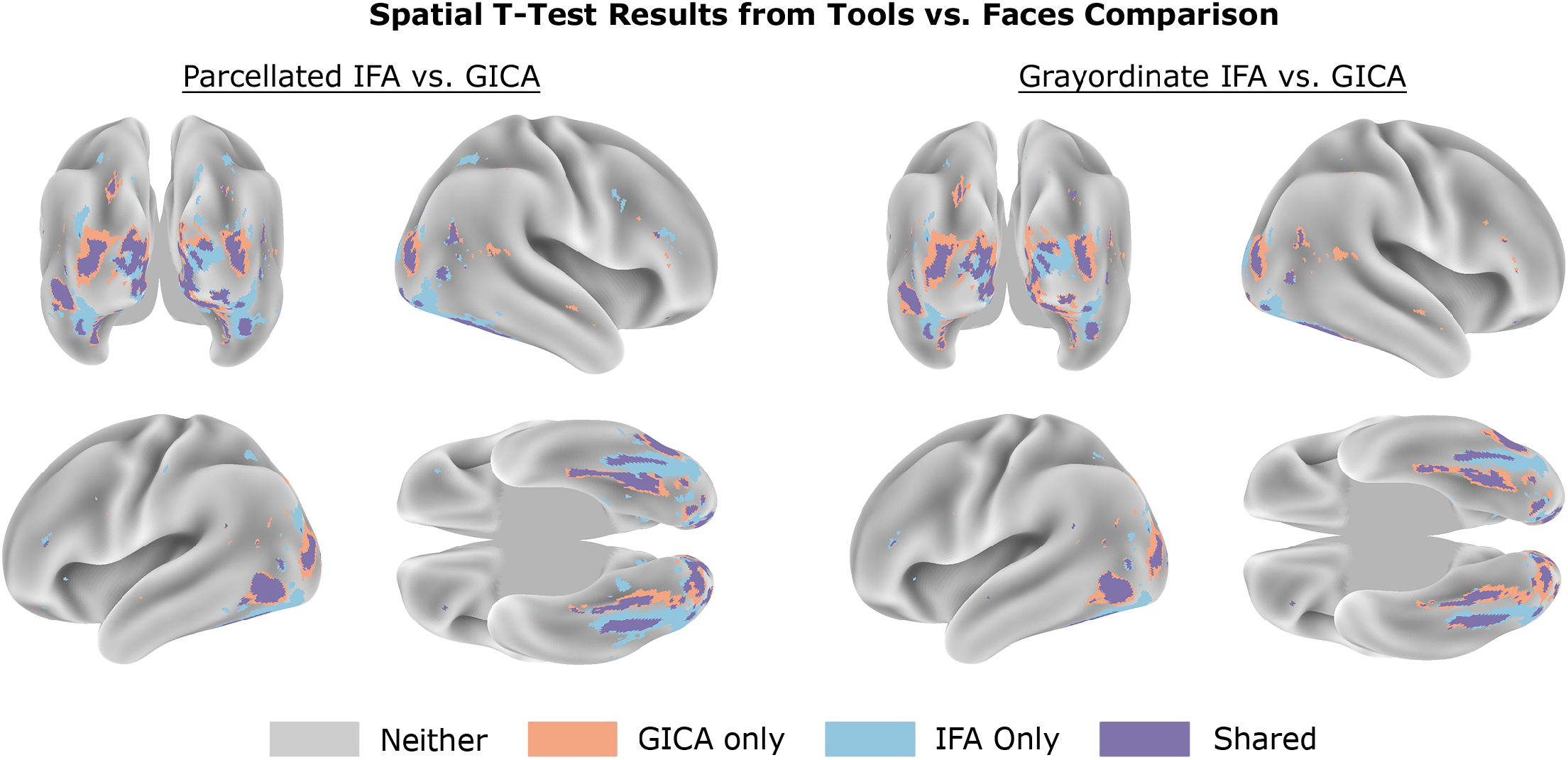
Spatial *t*-test differences between GICA and IFA pipelines for Face vs. Tools. Within each fold, paired vertex-wise *t*-tests were conducted between face and tool conditions for every spatial map produced by a pipeline. The resulting *p*-value maps were then collapsed into a single map by taking the minimum *p*-value at each vertex. IFA uniquely identifies more focal, task-relevant regions (e.g., fusiform face area), while GICA’s unique differences tend to reflect broader, less specific coverage in early visual areas already detected by both methods.

These comparisons reveal that the IFA pipelines more consistently and precisely detected task-specific regions. For both within-task comparisons, most of GICA’s unique significant differences were concentrated in early visual areas and tended to extend shared regions that both IFA and GICA identified, rather than task-relevant networks not captured by IFA. Conversely, IFA highlighted clusters in well-established category-selective parcels that GICA largely missed. In Face vs. Tools, IFA identified over twice as many significant vertices in the fusiform face area (FFA) (Kanwisher & Yovel, 2006), and almost uniquely captured both the right and left posterior inferior temporal (PIT) regions (Lafer-Sousa & Conway, 2013). For Relational vs. Match, all pipelines primarily detected early visual cortex differences, consistent with previous univariate analyses of these conditions within this task (Barch et al., 2013).

Applying 2D-LDA increased the number of significant *t*-test vertices across all pipelines by concentrating the discriminant signal into fewer components. GICA showed the largest gain, while IFA saw smaller improvements due to closer initial alignment to the discriminant spatial directions. These effects are reflected in the increased parcel coverage shown in Figure G, with the differences between IFA and ICA still present but reduced post-rotation.

### 3.3 Effect of Model Order on GICA vs. IFA Performance

The relative performance of IFA compared to GICA depends on both the selected model order and the location of discriminant signals within the eigenspectrum. As shown in Figure 5, IFA outperformed GICA across all model orders, effectively serving as an upper bound on GICA’s accuracy. The largest improvements appeared at lower model orders, where GICA’s eigenbasis contained little to no discriminant information. Consistent with the previous sections, the benefit of IFA was more pronounced for the Face vs. Tools and Relational vs. Match, where more stable filters were able to be identified.

**Figure 5:**
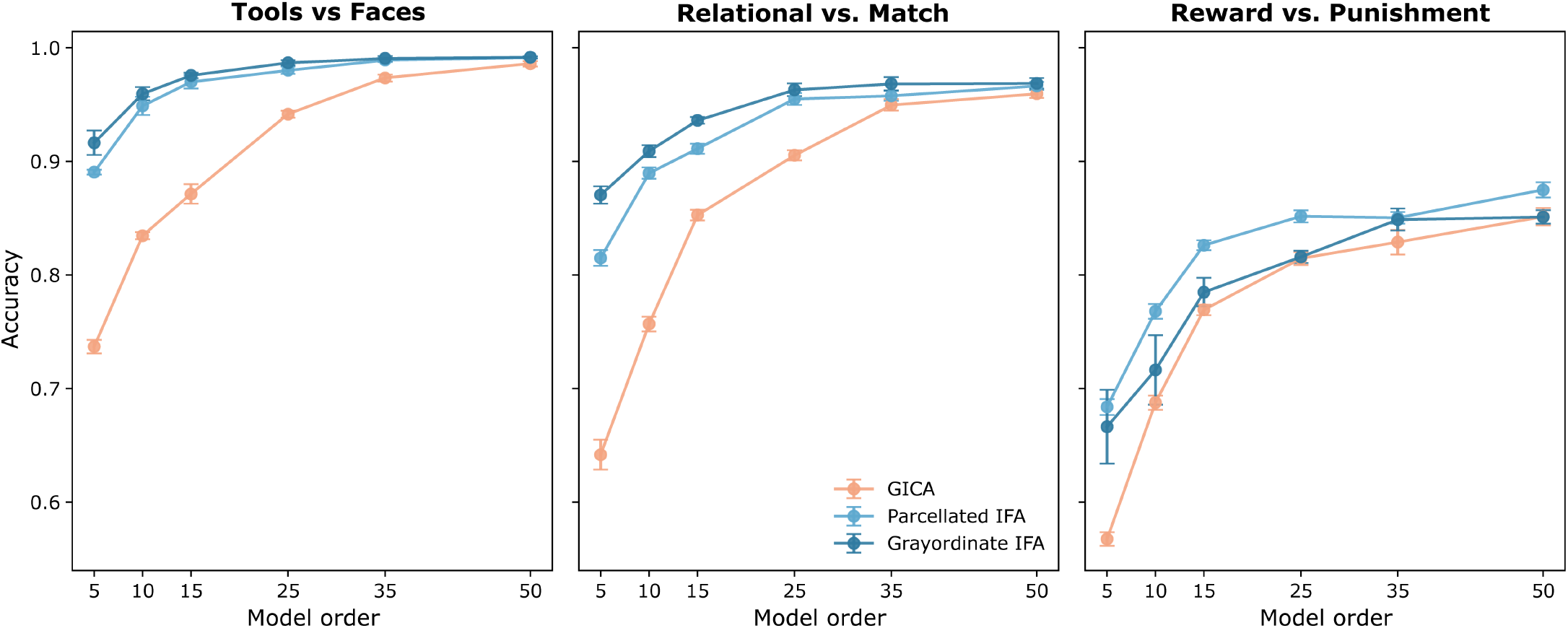
Effect of Model Order on Network Matrix Classification Accuracy.

For the Reward vs. Punishment task, IFA still outperformed GICA, but the advantage was smaller. Higher model orders were required to approach peak classification accuracy, likely due to the noisier and more complex group differences. Had additional discriminant filters per group been included, the lower model orders would likely have approached ceiling performance quicker. As in previous sections, Grayordinate IFA performed slightly better for Face vs. Tools and Relational vs. Match but was less robust on the Reward vs. Punishment task compared to Parcellated IFA.

While IFA consistently retained discriminant signals, GICA required substantially more components to approach comparable performance, reflecting its sensitivity to subtle effects as a function of model order and the placement of discriminant information within the eigenspectrum. Unlike GICA, IFA models discriminant structure explicitly rather than captured implicitly through model order selection. This approach reduces dependence on arbitrary model order choices, yielding more stable and reproducible solutions across datasets.

### 3.4 Simulated Data

We report the network matrix classification accuracy and the top discriminant spatial map accuracy for all pipelines applied to the simulated data in Table 3. In both datasets, where group-discriminative information was absent from the eigenspace retained by GICA, GICA failed to achieve above chance-level prediction for both network matrices and spatial maps. Parcellated IFA and Grayordinate IFA performed similarly on Dataset 1, where class membership in the discriminant space was well defined.

**Table 3:**
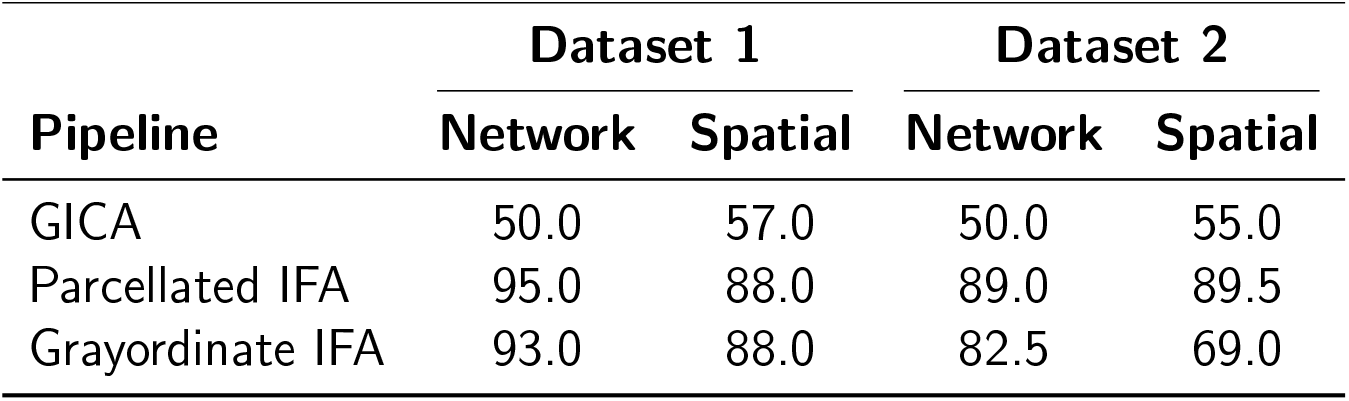
Simulated Data Classification Results for Network Matrices and Spatial Maps.

On Dataset 2, Parcellated IFA outperformed Grayordinate IFA in both domains. As established by Xu et al., 2019 when linking Common Spatial Patterns (CSP) to TSSF, Grayordinate IFA can be viewed as a special case of Parcellated IFA, where LDA is used as the tangent space classifier and class covariances are assumed to be **I**_*n*_^2^. Therefore, this method is Bayes-optimal only when class covariances in the discriminant subspace are identical across groups. Since this was not the case in Dataset 2, the SVM-based Parcellated IFA achieved better performance.

## 4 Discussion

We developed IFA to preserve class-specific information across all stages of group ICA. To achieve this, IFA extracts discriminant components through supervised dimensionality reduction, then applies PCA in the residual subspace to capture remaining variance. Starting with supervised dimensionality reduction ensures class-relevant information is retained regardless of the model order of the variance-explaining basis. Recovering the remaining variance-explaining subspace allows discriminant information to be embedded within biologically meaningful networks while remaining separable from structured noise. The resulting combined subspace is then unmixed using unsupervised FastICA, preserving spatial independence and supporting biological interpretability. Additionally, a post-hoc 2D-LDA projection provides a complementary view of how group differences are distributed across functional networks.

Across all analyses, IFA consistently improved sensitivity to group differences. This effect was most significant when between-group effect sizes are subtle and discriminant information is near-orthogonal to the GICA basis. In HCP task fMRI data, where GICA partially captured group differences, IFA still showed significant improvements in both the network matrix and spatial map domains. GICA tended to emphasize differences in shared networks, whereas IFA identified condition-specific patterns. When group differences dominate variance or enough components are retained, IFA offers limited additional benefit over GICA. In scenarios with noisy or ambiguous class labels, IFA is ultimately constrained by the amount of true discriminant signal present, as seen in the Reward vs. Punishment comparison.

Both IFA implementations achieved similar performance and consistently outperformed GICA. Parcellated IFA was more robust to within class variability due to TSSF’s flexibility in modeling between-subject variance. Grayordinate IFA allowed greater spatial flexibility and avoided dependence on any specific parcellation. However, it was less stable in noisy settings due to the use of large, ill-conditioned matrices and inability to model heterogeneity within groups. Additionally, because LOBPCG is an iterative solver, it can increase runtime by several hours when run on CPUs. With an appropriate preconditioner and execution on a GPU, runtime can be reduced to minutes, making it comparable to Parcellated IFA.

Incorporating discriminant components in place of purely variance-explaining components introduces a potential trade-off between enhancing group separation and preserving subject-level reconstruction accuracy. To quantify this, we compared reconstruction accuracy across pipelines (Figure I). GICA achieved marginally higher reconstruction accuracy, but this improvement offered no benefit for downstream analyses. In contrast, IFA’s discriminant components directly captured class-relevant effects. In practical applications where model order is not fixed for direct comparisons, the PCA step in IFA can capture the main sources of variance. The discriminant components computed from the outset ensure that class-specific information remains preserved, effectively balancing reconstruction and classification accuracy.

Embedding discriminant information into the group-level template contrasts with conventional unsupervised fMRI analysis methods. Historically, group-level approaches facilitate comparisons by constraining subject variability within a shared spatial subspace (Calhoun et al., 2001; Farahibozorg et al., 2021; Nickerson et al., 2017; Salman et al., 2019). Group differences are then only captured implicitly rather than modeled directly. Zhao et al. (2022) proposed Group LNGCA to overcome the limitation of variance-based dimensionality reduction. Rather than maximizing variance, this approach identifies a subspace that maximizes non-Gaussianity. Group LNGCA improved detection of group effects, but the unsupervised dimensionality reduction made performance sensitive to the number of non-Gaussian signals retained.

While some supervised ICA variants have attempted to address this limitation, most introduced class labels only during unmixing (Du et al., 2023; Maneshi et al., 2016; Tabas et al., 2014). Sui et al., 2009 introduced an early exception to this, where they incorporated class information in both the dimensionality reduction and unmixing. However, in addition to being limited to task-derived features, the method relied on a post-hoc ranking of PCA components based on a group separation criterion. Since the initial decomposition prioritizes variance rather than discrimination, the reranked components are not guaranteed to align with directions that capture subtle differences between groups.

IFA directly addresses these shortcomings for two-class discrimination. An important next step will be extending the supervised dimensionality reduction step to support multiclass classification and regression. To support diagnostic or clinical applications, further refinements will be needed to improve consistency and interpretability. These include exploring alternative regularization strategies, such as ordered weighted *ℓ*_1_ penalties (Bondell & Reich, 2008; Figueiredo & Nowak, 2016), which promote structured sparsity and group-level feature selection. Additionally, hyperparameter tuning procedures may need to be refined to better align with specific hypotheses, rather than relying solely on general reconstruction metrics like *R*^2^.

Independent Filter Analysis provides a framework for modeling both shared and subgroup-specific variance throughout a group-level modeling pipeline. This dual focus supports both exploratory and hypothesis-driven analyses in neuroimaging studies targeting subtle phenotypic differences or early diagnostic markers.

## 5 Data and Code Availability

The code used for implementing and testing IFA is publicly available on GitHub: https://github.com/zainsouwei/IFA.git.

Data were provided by the Human Connectome Project, WU-Minn Consortium (Principal Investigators: David Van Essen and Kamil Ugurbil; 1U54MH091657) funded by the 16 NIH Institutes and Centers that support the NIH Blueprint for Neuroscience Research; and by the McDonnell Center for Systems Neuroscience at Washington University.

## 6 Author Contributions

**Zain Souweidane**: Conceptualization, Formal analysis, Methodology, Software, Validation, Visualization, Writing – original draft, Writing – review & editing. **Alberto Llera**: Conceptualization, Methodology, Writing – review & editing. **Stephen M. Smith**: Conceptualization, Formal analysis, Funding acquisition, Methodology, Supervision, Validation, Writing – review & editing. **Christian F. Beckmann**: Conceptualization, Formal analysis, Funding acquisition, Methodology, Resources, Supervision, Validation, Writing – review & editing.

## A Dual Regression Hyperparameter Selection

The ElasticNet optimization is formulated as

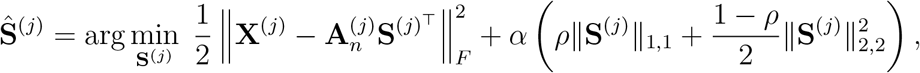

where for a given subject *j*, **X**^(*j*)^ ∈ ℝ^*t*×*s*^ is their fMRI data, 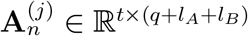 the matrix of normalized component time courses from the first stage of dual regression, and 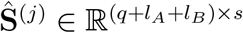 are the estimated spatial maps from the second stage of dual regression (with ∥ · ∥_1,1_ and ∥ · ∥_2,2_ denoting entrywise *ℓ*_*p,q*_ norms, where ∥ · ∥_2,2_ = ∥ · ∥_*F*_).

Ideally, the optimal ElasticNet hyperparameters (*α, ρ*) maximize reconstruction across all training subjects. However, doing so would require *m* · *n* dual regressions, where *n* is the number of subjects and *m* the number of hyperparameter configurations. To efficiently estimate the hyperparameters, a subset of training subjects is selected using stratified random sampling with shuffling to maintain the two-class balance. *α* ∈ [10^−3^, 10] was sampled in log space while *ρ* ∈ [*ϵ*, 1 − *ϵ*], with *ϵ* = 10^−4^, was sampled linearly. We set *ϵ* = 10^−4^ to prevent the hyperparameter search from evaluating degenerate edge cases where the ElasticNet collapses to a purely *ℓ*_1,1_ or *ℓ*_2,2_ penalty, which can lead to instability during model fitting. For each candidate hyperparameter configuration, selected using Gaussian Process Optimization, the following procedure is performed:

1. For each subject in the subsample, time points are split into training and held-out sets.
2. ElasticNet is fit on the training data to estimate spatial maps.
3. The R^2^ score is computed on held-out data for that subject.
4. The average R^2^ across all subjects defines the configuration’s performance.
5. Steps 1–4 repeat until convergence.

This was implemented with `sklearn.linear model.ElasticNet`, which fits an independent regression per grayordinate. This corresponds to an *ℓ*_1,1_ + *ℓ*_2,2_ penalty on the full coefficient matrix and should not be confused with joint-sparsity models such as `MultiTaskElasticNet`, which imposes an *ℓ*_2,1_ penalty.

## B Discriminant Spatial Maps

The spatial maps obtained from the unmixed combined subspace are not guaranteed to align with the spatial patterns in the subspace that maximally separate the groups. We apply 2D-LDA (Ye et al., 2004) on subject-level spatial maps to find the linear combinations that best separate the groups. A derivation is provided to illustrate how this transformation can be interpreted as a rotation of the spatial maps that enhances between-group separability.

Let 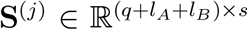 denote the spatial maps for subject *j*, and let {A, B} be the two groups with *n*_*A*_ and *n*_*B*_ subjects, respectively.

Define the group-mean spatial maps as:

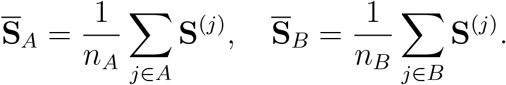

We seek a projection direction 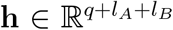 that maximizes the Euclidean distance between projected group means:

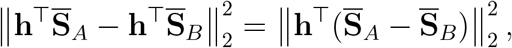

normalized by within-group variability (i.e., Mahalanobis distance). Formally, we solve:

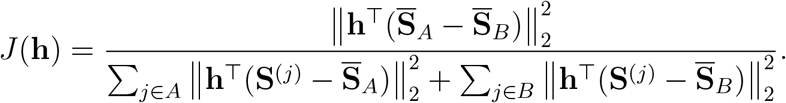

Rewriting this as a generalized Rayleigh quotient:

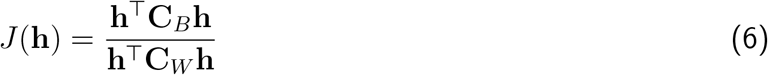

where

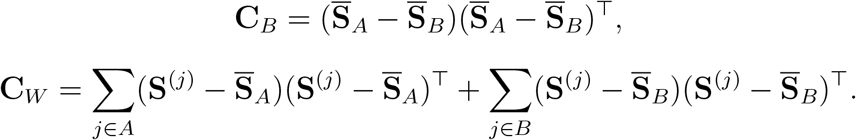

The solution to Equation 6 can be efficiently found by solving the generalized eigenvalue problem:

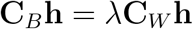

which yields a **C**_*W*_ orthogonal matrix 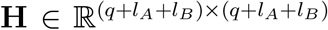. We transform each subject’s spatial maps by:

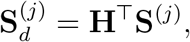

such that the top rows of 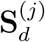 capture the directions of maximal between-group separation, while the lower rows capture the least.

To improve stability in high-dimensional settings, we regularize **C**_*W*_ using the Oracle Approximating Shrinkage (OAS) estimator.

## C SPADE and TSSF Background

SPADE identifies spatial directions that maximize the variance explained for one group while minimizing it for the other, as shown in Figure 6. The generalized eigenvalue problem used by SPADE (Equation 5) can be re-expressed as a standard eigenvalue decomposition by whitening both sides by 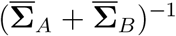 (Fukunaga & Koontz, 1970), yielding:

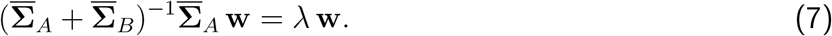

**Figure 6:**
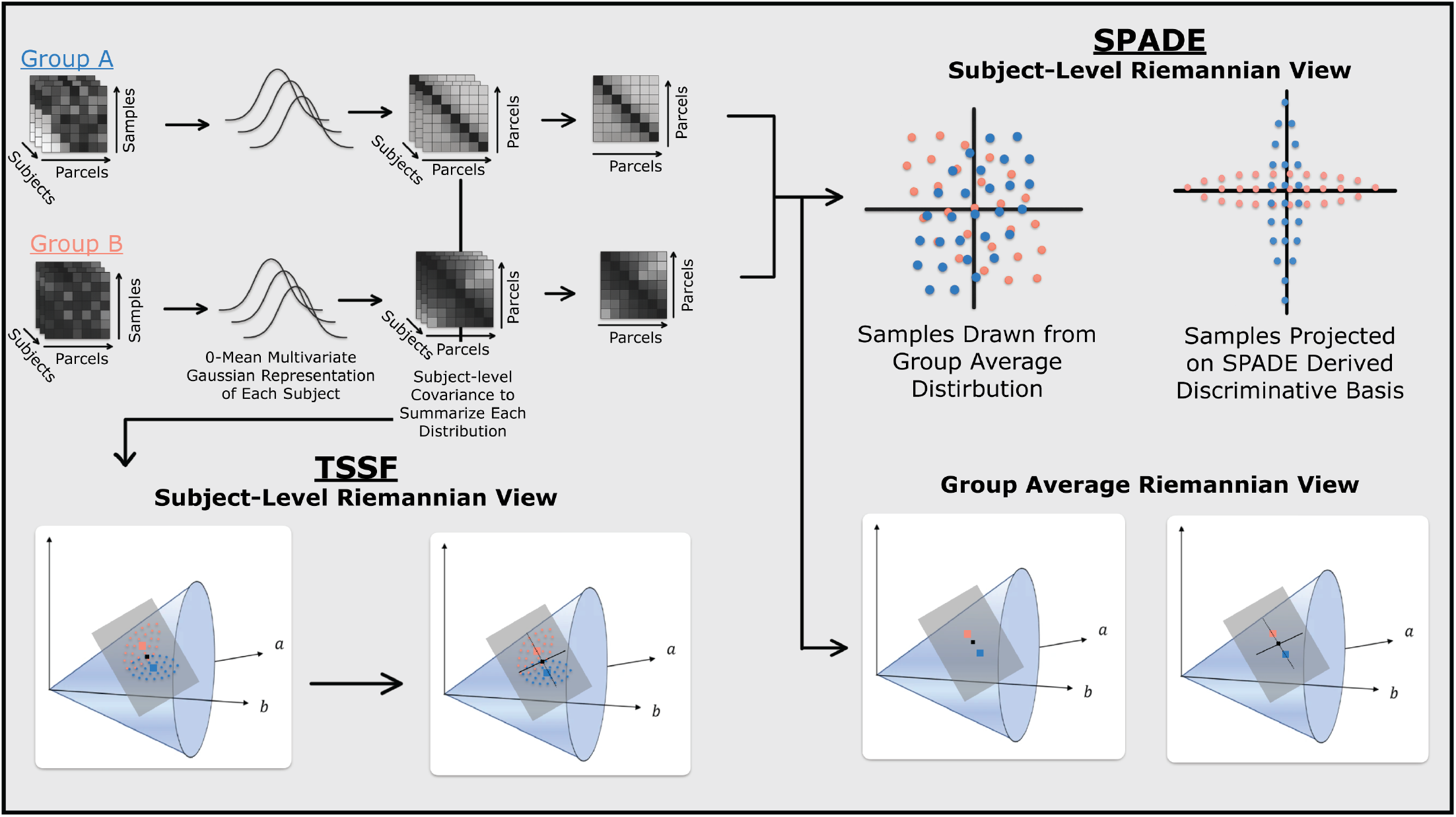
Conceptual overview of SPADE vs. TSSF for modeling group differences in fMRI. Each subject’s time series is modeled as samples from a zero-mean multivariate Gaussian, summarized by a subject-specific covariance matrix. These subject covariances are averaged to define a class-level distribution. *SPADE (Right):* (Top) Identifies spatial filters that, on average, maximize variance for one group’s class-average covariance while minimizing it for the other. This corresponds to finding directions that optimally separate samples drawn from the group-average distributions. (Bottom Right) In the Riemannian framework, this is equivalent to finding directions that maximize the distance between class average covariances. *TSSF (Bottom Left):* Operates at the subject level by treating each subject’s covariance as a point on the Riemannian manifold of symmetric positive definite matrices. These points are projected to the tangent space at a shared reference. This approach models between-group differences and within-group variability and enables flexible linear classification in a locally linearized space.

**Figure 7:**
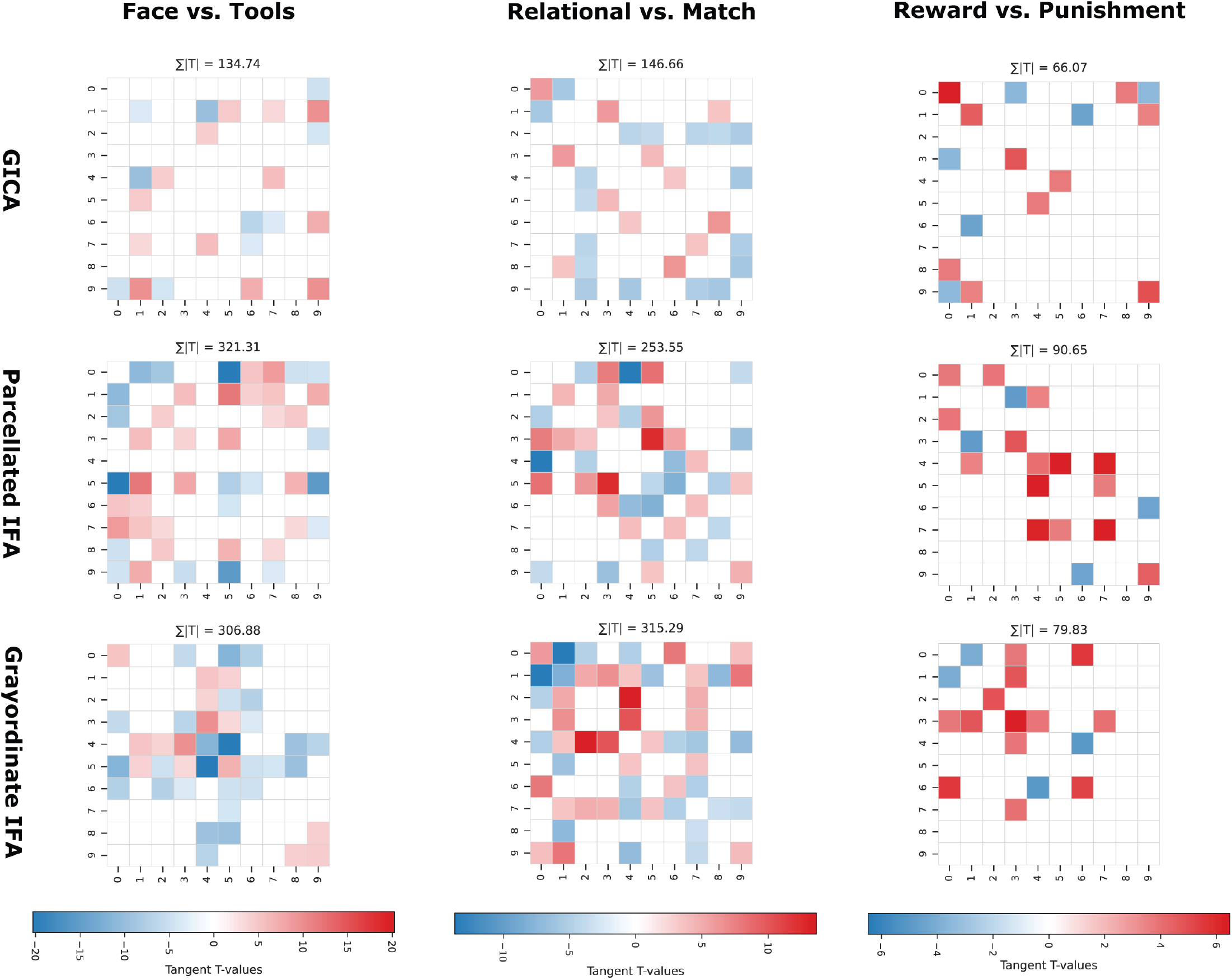
Group Differences in Tangent-Space Network Matrices. Heatmaps show statistically significant *t*-values from permutation-based t-tests (max-statistic correction, *α* = 0.05) applied to tangent-space network matrices. Non-significant connections were thresholded to zero, and results were unvectorized to symmetric matrices for visualization. Red indicates connections with significantly higher values in Group A, blue indicates higher values in Group B, and white indicates no significant difference. Each column represents a task (Face vs. Tools, Relational, Incentive), and each row corresponds to a pipeline (GICA, IFA Parcellated, IFA Grayordinate). The sum of absolute *t*-values (∑|*T* |) is reported in each heatmap to quantify overall discriminative effect strength.

The resulting eigenvectors define orthogonal directions in this shared space (i.e., 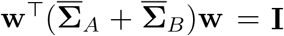), with each eigenvalue *λ* indicating the proportion of variance contributed by Group A along the corresponding direction. The remaining variance, 1 − *λ*, is contributed by Group B. The filters then represent the directions of maximal variation from the shared reference. From a geometric perspective, this is conceptually similar to identifying directions on the Riemannian manifold of symmetric positive definite matrices that maximize the distance between the class-average covariances (Barachant et al., 2010).

TSSF builds on this intuition by incorporating subject-level variability, rather than only comparing group averages. TSSF whitens each subject’s covariance matrix using the shared covariance, projecting them into a common vector space where each subject is represented as a displacement from a shared reference. This vector space is theoretically equivalent to the plane tangent to the average (i.e., Fréchet mean) of all subject covariances on the manifold. A linear classifier is trained in this space to identify the direction that best separates the classes, and an eigenvalue decomposition is performed on the classifier’s weight matrix to derive the spatial filters. By leveraging Riemannian geometry and focusing on between-subject variance, TSSF identifies more robust discriminant features, particularly when subject-level heterogeneity is high. The relationship between SPADE and TSSF is illustrated in Figure 6.

This process starts by selecting a Riemannian metric and computing the Fréchet mean, defined as 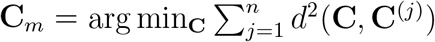, across all subject covariance matrices 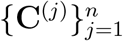. Each subject’s covariance is then projected to the tangent space at **C**_*m*_ using the logarithmic map 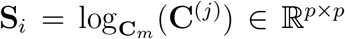, where **S**^(*j*)^ is the symmetric matrix in the tangent space. The upper triangular of **S**^(*j*)^ is vectorized into 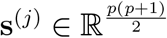, with off-diagonal elements scaled by 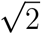. These vectors are used to train a linear classifier to separate groups. The learned weight vector 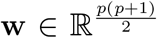 is reshaped back into a symmetric matrix **S**_*w*_ ∈ ℝ^*p*×*p*^ in the tangent space. An EVD is then performed on **S**_*w*_:

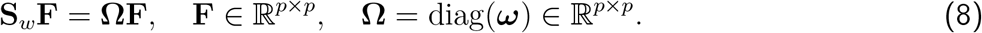

The eigenvectors **F** form an orthonormal basis that reconstructs the line normal to the tangent classifier’s decision boundary. The magnitude of an eigenvalue,***ω***, indicates the contribution of its associated eigenvector to supporting the Riemannian distance between classes (Barachant et al., 2010). The top and bottom eigenvectors are chosen as the spatial filters **F**_*g*_, which define directions that most strongly push the positive and negative classes, respectively, away from the shared reference point.

The primary method described in the original TSSF formulation involves mapping the symmetric weight matrix **S**_*w*_ to the manifold via the exponential map to form 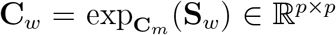. Then a GEVD is applied:

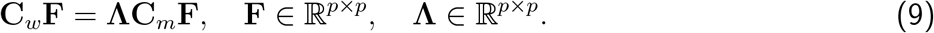

As established in Appendix A of Xu et al., 2019, the eigenvectors derived from Equation 8 and Equation 9 are equivalent, while the eigenvalues differ by a logarithmic transformation induced by the Affine Invariant Riemannian Metric (AIRM). The GEVD in Equation 9 is inherently linked to the AIRM (Barachant et al., 2010), so the alternative use of Equation 8 preserves the eigenvalue magnitudes under the selected metric and avoids enforcing a specific geometric structure, allowing greater flexibility across metric choices. Our analysis revealed minimal differences between the two approaches in terms of the derived spatial patterns.

## D Projection of Parcellated Filters to Grayordinate Space

We use a dual regression approach to project parcellated discriminant filters 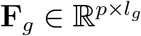 to grayordinate space. This is applied separately for each group *g* ∈ {A, B}.

Let 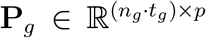 denote the temporally concatenated parcellated data for all *n*_*g*_ subjects with *t*_*g*_ samples in group *g*, and 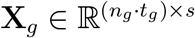 the corresponding grayordinate data.

First, we compute the group-level projection of the parcellated data onto the filter space:

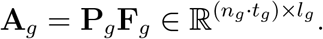

We then regress the grayordinate data onto this projection:

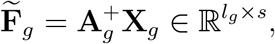

where 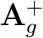 denotes the pseudoinverse of **A**_*g*_.

Since **X**_*g*_ is high-dimensional, this regression is performed efficiently using a block matrix formulation. 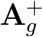 can be partitioned into subject-specific blocks:

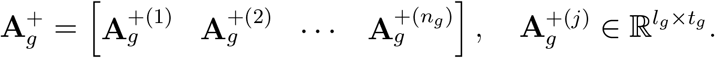

Note each 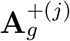 is not the pseudoinverse of subject *j*’s data, but rather the portion of the overall group-level pseudoinverse 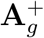 that corresponds to subject *j*’s data block. Similarly, we decompose the group-level, grayordinate data:

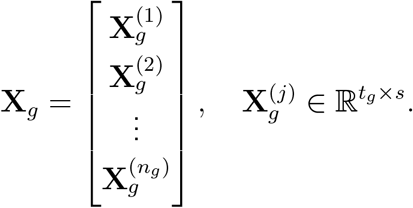

The final transformation is expressed as the product:

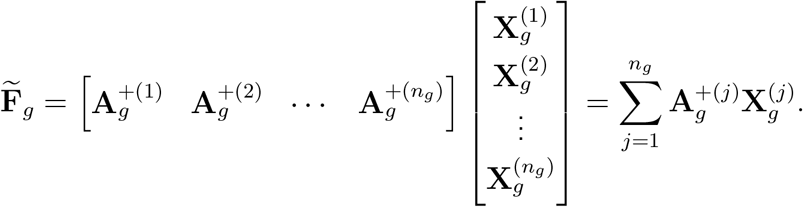

This formulation performs a partial regression per subject and combines the results across the group, enabling memory-efficient parallel implementation (Golub & Loan, 2013).

## E Log-Euclidean Grayordinate SPADE

The log-Euclidean metric can additionally be used in the grayordinate-based implementation. Since the symmetric positive definite (SPD) manifold with the Log–Euclidean metric is globally isometric to the Euclidean space of symmetric matrices (Arsigny et al., 2006, 2007), all distances, geodesics and Fréchet means reduce to ordinary Euclidean computations in the log-domain. In practice, only the matrix logarithm is needed to project each covariance onto the tangent space. As shown below, this can be efficiently computed via a Singular Value Decomposition (SVD).

The group average can be formed directly in the log space. For each subject *j* in group *g* (with *g* ∈ {*A, B*}) and data matrix 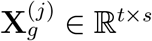, we compute their covariance matrix, 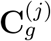, via the SVD. First, we decompose 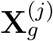 as

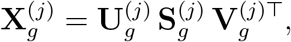

where 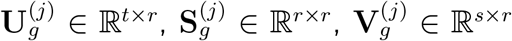, and 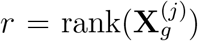. Since the covariance structure is related to the squared singular values, we define the subject *j*’s covariance matrix as

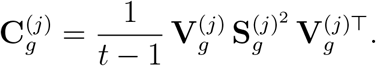

Taking the matrix logarithm yields

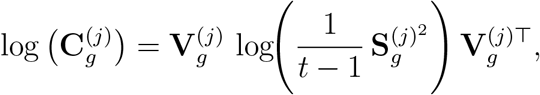

where 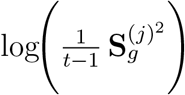 is computed element-wise on the diagonal.

The log-Euclidean group average is then obtained by averaging the log-covariance matrices across subjects:

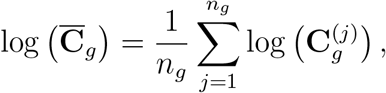

with *n*_*g*_ being the number of subjects in group *g*. This average in the log domain provides a robust estimate of the group covariance under the Log-Euclidean metric.

Once the log-domain covariances for groups A and B have been computed, the discriminative filters are extracted by solving the eigenvalue problem:

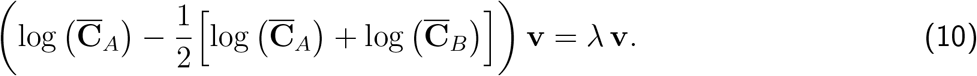

and

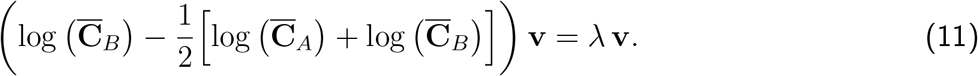

In this formulation, 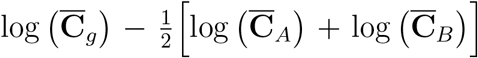 represents the deviation of group *g*’s log-covariance matrix from the midpoint between groups A and B. The top eigenvectors obtained from solving both equations Equation 10 and Equation 11, using LOBPCG correspond to the group discriminant filters. The filters are then converted into signal-space activation maps via the Haufe-transform (Section 2.2) using their respective group-average MIGP-derived covariances.

## F Network Matrix T-Test Results

## G Spatial Map T-Test Results

These figures show parcel-level summaries of vertex-wise *t*-test results for ICA- and IFA-derived representations. For each MMP parcel, we calculated the percentage of significant vertices that were unique to IFA (blue), unique to GICA (orange), or shared between methods (purple). Only parcels with more than 5% total significant coverage are displayed. Significance was determined using permutation-based vertex-wise *t*-tests with maximum-statistic correction (*α* = 0.05, 10,000 permutations). To generate summary maps, we retained the minimum *p*-value across components at each vertex. Analyses were also conducted on the 2D-LDA–rotated spatial maps. Basic refers to spatial maps obtained directly from ICA and dual regression, while Discriminant refers to the 2D-LDA rotated subject-level maps.

### G.1 Working Memory Task (Face vs. Tools)

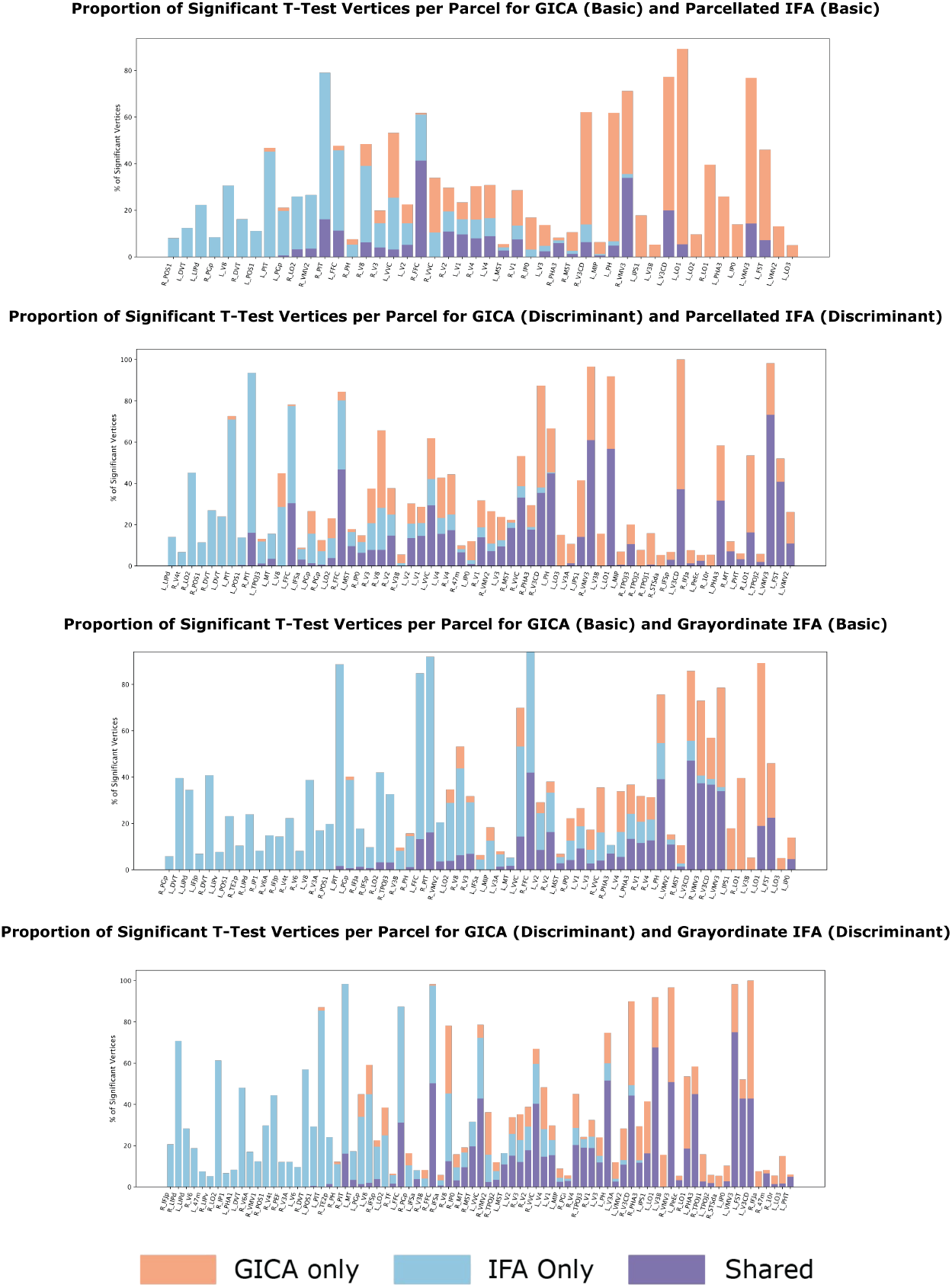

### G.2 Relational Task (Relational vs. Match)

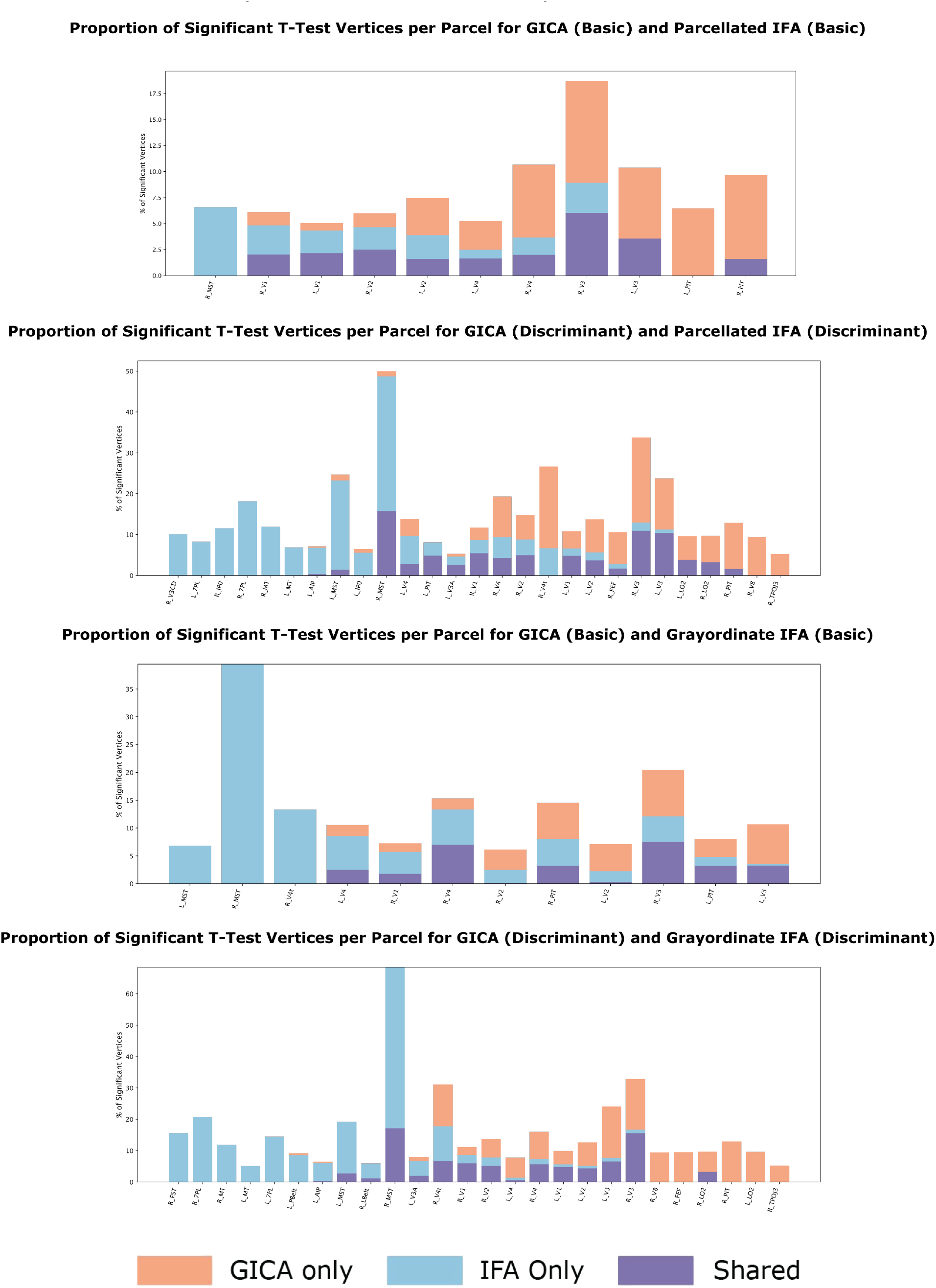

## H Simulated Discriminant Subspace

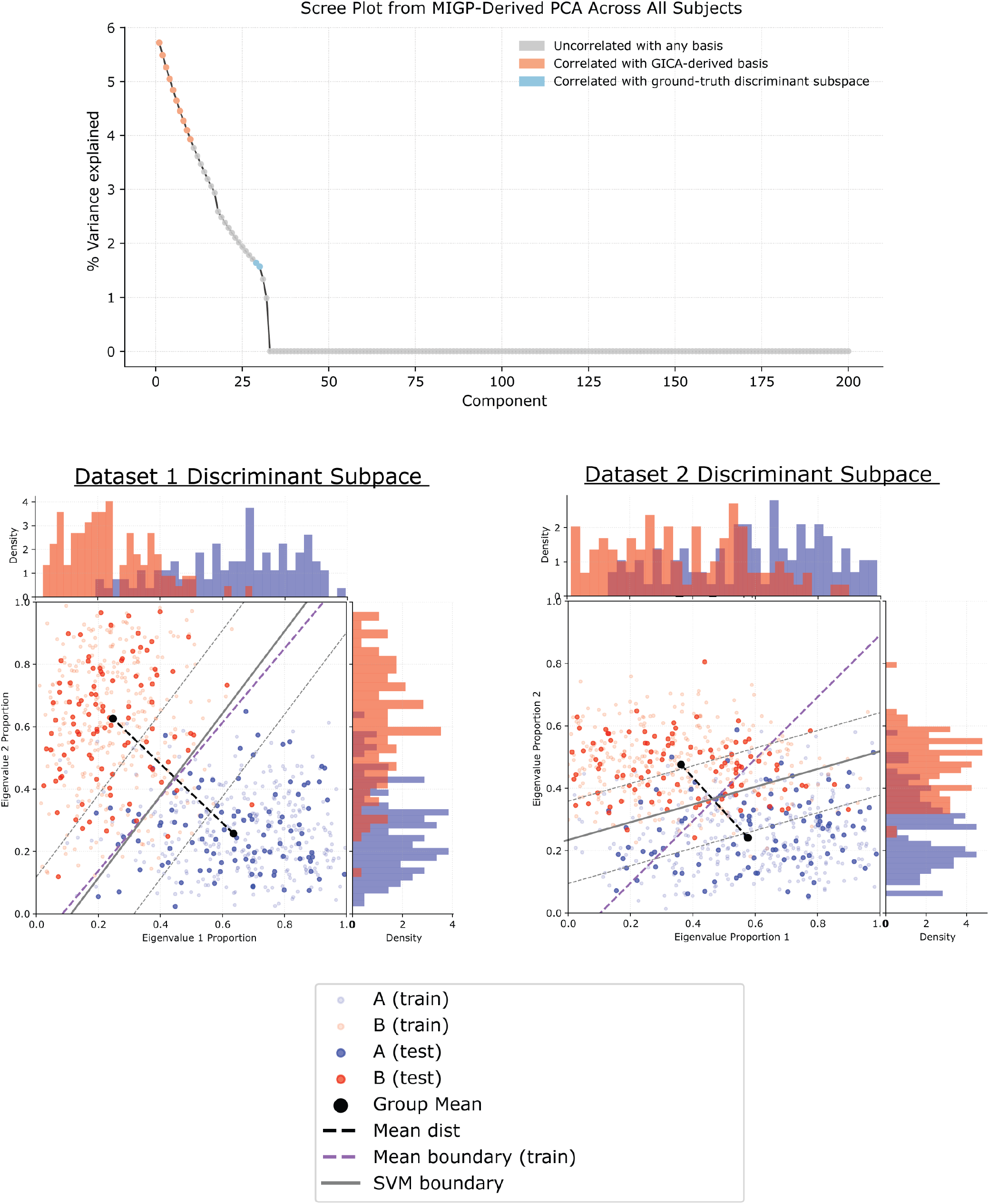

(**Top**) Scree plot from MIGP-derived PCA across all subjects. Components correlated with the GICA-selected basis are highlighted in orange, those correlated with the ground-truth discriminant subspace in blue, and the remaining components are uncorrelated (gray). It can be seen that the entire discriminant subspace was left out of the final GICA subspace. (**Bottom**) Two-dimensional discriminant subspaces for Dataset 1 (left) and Dataset 2 (right). Each point corresponds to a subject’s projection onto the two discriminant eigenvectors. Group means, mean boundaries, and SVM decision boundaries (solid gray) are indicated to visualize the linear separability of the classes. Dataset 1 shows classes that are well represented by their class-specific eigenvalues, while Dataset 2 shows one noisier class.

## I Reconstruction Accuracy

**Figure 9:**
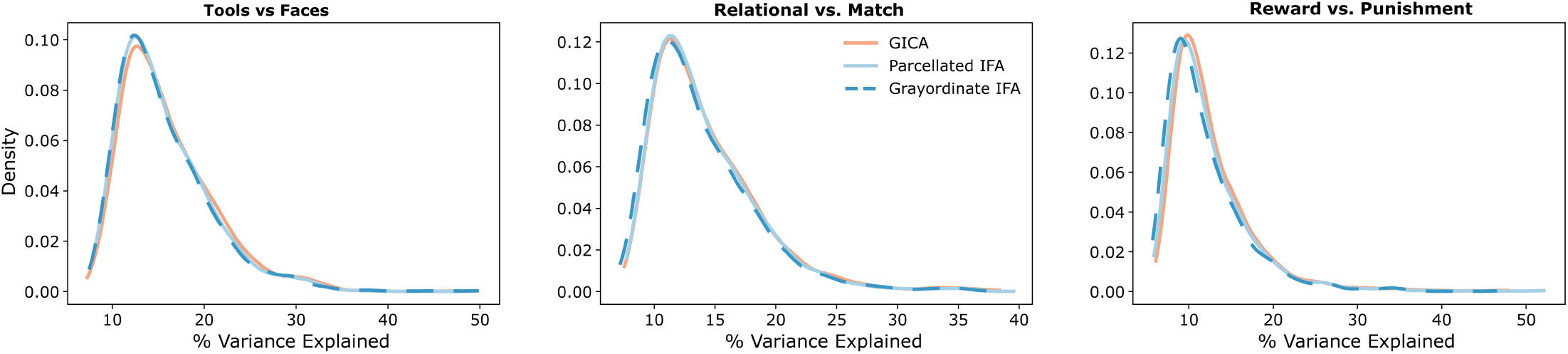
Kernel density estimates of subject-level reconstruction accuracy across five folds for each model and task, reflecting the percentage of variance in each subject’s original data explained by reconstructions using their dual-regression derived spatial maps and time series.

## Footnotes

1 We additionally evaluated results from computing filters in the full (non-residual) space, as well as swapping the order and computing filters in the residual space of the variance-explaining subspace. While the effects of the ordering are dependent on the contrast ratio of discriminant information and variance explaining information, we found computing the discriminant basis first to be the most robust across different conditions and model orders. Future work will explore integrated approaches that jointly optimize the full signal space.

2 This is a high-level equivalence that overlooks the practical tradeoff of performing discriminant analysis in parcellated space versus grayordinate space.

